# Critical assessment of approaches for molecular docking to elucidate associations of HLA alleles with Adverse Drug Reactions

**DOI:** 10.1101/296574

**Authors:** Kerry A Ramsbottom, Dan Carr, Andrew R Jones, Daniel J Rigden

## Abstract

Adverse drug reactions have been linked with genetic polymorphisms in HLA genes in numerous different studies. HLA proteins have an essential role in the presentation of self and non-self peptides, as part of the adaptive immune response. Amongst the associated drugs-allele combinations, anti-HIV drug Abacavir has been shown to be associated with the HLA-B*57:01 allele, and anti-epilepsy drug Carbamazepine with B*15:02, in both cases likely following the altered peptide repertoire model of interaction. Under this model, the drug binds directly to the antigen presentation region, causing different self peptides to be presented, which trigger an unwanted immune response. There is growing interest in searching for evidence supporting this model for other ADRs using bioinformatics techniques. In this study, *in silico* docking was used to assess the utility and reliability of well-known docking programs when addressing these challenging HLA-drug situations. Four docking programs: SwissDock, ROSIE, AutoDock Vina and AutoDockFR, were used to investigate if each software could accurately dock the Abacavir back into the crystal structure for the protein arising from the known risk allele, and if they were able to distinguish between the HLA-associated and non-HLA-associated (control) alleles. The impact of using homology models on the docking performance and how using different parameters such as including receptor flexibility affected the docking performance, were also investigated to simulate the approach where a crystal structure for a given HLA allele may be unavailable. The programs that were best able to predict the binding position of Abacavir were then used to recreate the docking seen for Carbamazepine with B*15:02 and controls alleles. It was found that the programmes investigated were sometimes able to correctly predict the binding mode of Abacavir with B*57:01 but not always. Each of the software packages that were assessed could predict the binding of Abacavir and Carbamazepine within the correct sub-pocket and, with the exception of ROSIE, was able to correctly distinguish between risk and control alleles. We found that docking to homology models could produce poorer quality predictions, especially when sequence differences impact the architecture of predicted binding pockets. Caution must therefore be used as inaccurate structures may lead to erroneous docking predictions. Incorporating receptor flexibility was found to negatively affect the docking performance for the examples investigated. Taken together, our findings help characterise the potential but also the limitations of computational prediction of drug-HLA interactions. These docking techniques should therefore always be used with care and alongside other methods of investigation, in order to be able to draw strong conclusions from the given results.

## 1. Introduction

An adverse drug reaction (ADR) is a harmful or unpleasant reaction, resulting from the use of medicinal products. Type A reactions are those that are dose-related. Idiosyncratic drug reactions (IDRs) or Type B hypersensitivity reactions are dose-independent, occurring in some but not all people [1]. The incidence of ADRs have increased globally from 2.2 million in 1994 to 10 million in 2014 [2]. This is therefore a very important issue which needs to be addressed.

These ADRs have been linked with specific Human Leukocyte Antigens (HLA) in numerous studies, whereby individuals carrying particular alleles of HLA genes are at higher risk of developing adverse reactions to particular drugs [3–5]. HLA gene products play a key role in the adaptive immune response, presenting peptides (self and non-self) to a T cell complex to elicit a response when needed. The HLA system is highly variable, both in individuals and in populations. Individuals carry multiple HLA genes with similar functions: A, B, C in class I, or DRA, DRB, DQA, DQB and others in class II. Class I gene products are responsible for presentation of peptides from pathogens internal to cells, such as viruses. Class II gene products present peptides from extracellular pathogens.

HLA alleles are given a unique identifier, following a detailed and well-established nomenclature system, such as ‘HLA-B*57:01’. The identifier always has the prefix HLA- and then the gene identifier (A, B, C for class I HLA genes, or DRA, DRB, DQA, DQB and others for class II HLA genes) followed by a “*” separator and a set of numbers separated into groups. The first two numbers after the ‘*’ separator give the allele group, originally defined by serotyping, and the next two numbers following ‘:’ are unique for the specific HLA protein sequence. Further sets of digits are possible after additional colon separators i) to identify alleles different at the exon (DNA)-level but causing no change to the protein sequence (synonymous substitutions), and then ii) for substitutions in intronic regions e.g. ‘HLA-B*57:01:01:01’. For consideration of HLA-ADRs, four digit resolution (i.e. resolved to the protein sequence level only) is generally considered sufficient [6].

The role of HLA in ADRs has been hypothesised in three main ways. The *Hapten* model predicts that the drug binds covalently to a self protein, and is processed via HLA molecules to the presented peptide; this drug-protein combination then being recognised as being non-self and initiating an immune response. The *Pharmacological Interaction (PI)* model predicts that the drug binds non-covalently, directly to the immune receptors; mainly T-cell receptors or HLA. The *Altered Peptide Repertoire* model states that the drug interacts with the HLA molecule directly and non-covalently, leading to a different self-peptide set being presented, which is recognised as foreign, and thus eliciting the immune reaction [7]. Illing *et al*. showed that the Abacavir modifies the anchor residue for the binding peptide in the F-pocket, altering the binding specificity for peptides in B*57:01 but not B*57:03 [8].

ADRs are associated with different HLA alleles for numerous different drugs. The ‘HLA and Adverse Drug Reactions’ database on the Allele Frequency Net Database website [9, 10] allows users to search for studies showing associations between different HLA alleles and ADRs. The current, most strongly associated ADR is that of Abacavir (an anti-retroviral drug) with HLA-B*57:01. If certain alleles have been significantly associated with ADRs, patients can be screened prior to being given the drug to predict if an ADR is likely to occur. Mallal *et al*. showed how screening for HLA-B*57:01 alleles can reduce the risk of hypersensitivity reactions in patients receiving Abacavir [11]. While there is still some disagreement which of the models best explains how they interact with drugs to cause ADRs, Illing *at al*. have demonstrated the Altered Peptide Repertoire model with high confidence for Abacavir, including a crystal structure of Abacavir bound to the antigen presenting region of HLA-B*57:01, as well as proteomics evidence for different peptides being presented than in the unbound case. As a result, many researchers investigating ADRs now work under the assumption that this hypothesis explains a high proportion of HLA ADRs observed, although much debate continues. There is therefore considerable and growing interest in searching for evidence supporting this mode for other ADRs using modelling and bioinformatics techniques, for example using *in silico* molecular docking [8, 12–15].

Molecular docking is used to predict the preferred orientation of a molecule when bound to another in a stable complex. Most docking programs assume the target to be rigid and allow ligand flexibility [16]. Protein-ligand docking can be used to aid understanding of biological processes and drug design [17, 18]. Docking gives a prediction of the structure of the ligand-receptor complex using computational methods by first sampling the conformations of the ligand in the active site and then ranking these conformations using a scoring function as a proxy for the free energy of interaction [19].

Molecular docking is being used increasingly commonly for investigating HLA-mediated ADRs [8, 12, 13, 15, 20–25]. The HLA structure presents unusual challenges for molecular docking protocols. HLAs bind peptides in a long hydrophobic cleft formed between the α-helices and β-sheet platform. This cleft is much larger than the naturally evolved binding sites that proteins have for small organic molecules. The polymorphic residues located along this cleft determine the size and stereochemistry of the subsites [26]. The peptide binding groove contains six subsites (Fig 1). The specificity of peptide binding is determined by the interactions between anchor residues on the peptide side chains and two or more of these subsites [27]. Therefore care must be taken when using docking methods to investigate these complex cases. The purpose of this exercise is to compare multiple docking programs to assess their performance on the challenging HLA-ADR cases.

**Fig 1.**
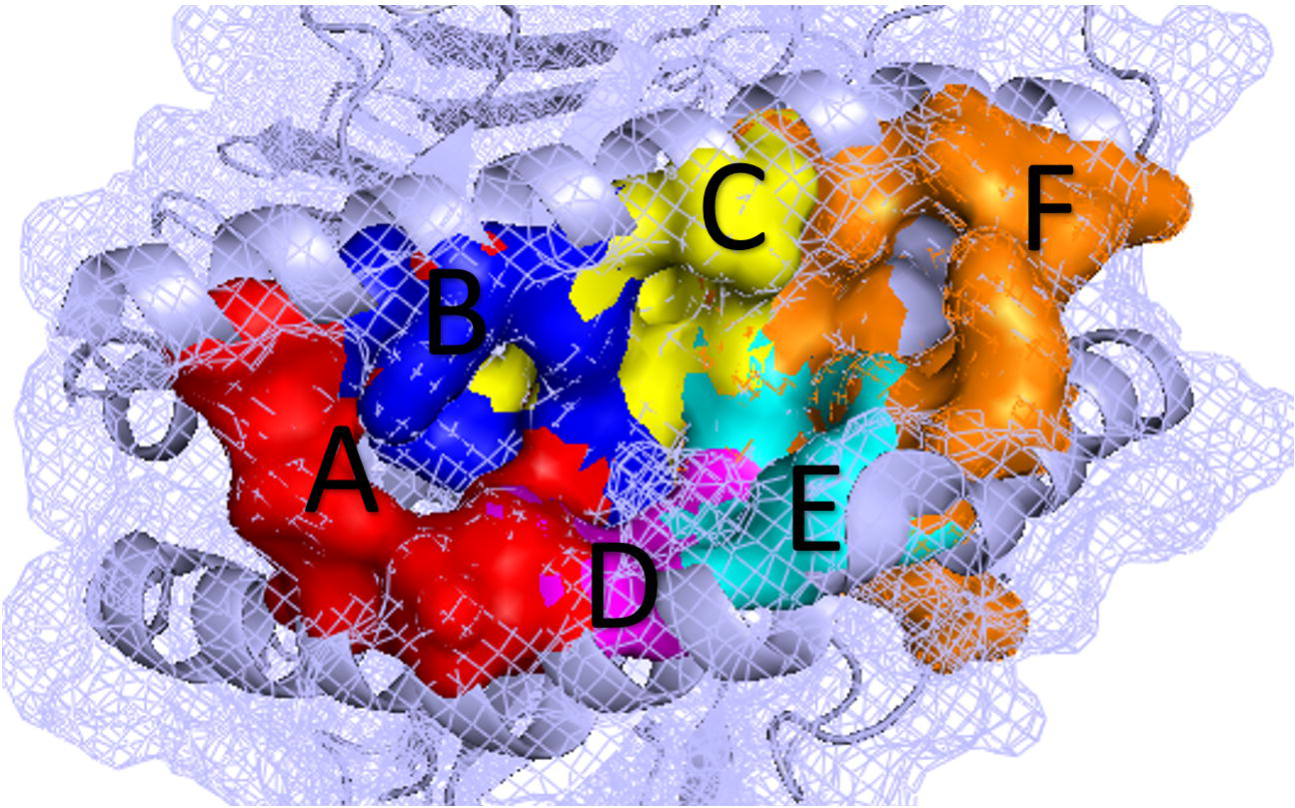
Organisation of the subsites along the HLA peptide binding groove. The peptide-binding groove of the HLA molecule is separated into 6 different pockets (A-F) [28, 29], as shown here. Image created using PyMOL [30].

Four different freely available and commonly used programs were used for docking the compounds with the target alleles – SwissDock, ROSIE, AutoDock Vina and AutoDockFR, as follows. The SwissDock [31] server is an online tool based on the EADock DSS [32] engine. Target and ligand structures can be automatically prepared for docking through the server. Target structures can be selected via PDB records, or user-defined structures can be uploaded in various supported formats. Ligands can be selected through the ZINC database or by uploading structure files. A range of docking parameters can be set, including docking type, enabling the user to select a desired docking time and exhaustiveness, and defining the search space [31]. Due to it being an online tool, it is very accessible and can be used without the technical knowledge required for some of the more complex software.

The Rosetta Online Server that Includes Everyone (ROSIE) is an online version of the Rosetta 3 software. The server includes different Rosetta protocols, including RosettaLigand which allows small molecules to be docked into proteins. The target structure must be provided in PDB format. For best results, residues that Rosetta does not natively recognise (e.g. waters, co-factors or metal ions) should be removed prior to submission. An SDF file containing the conformers of a single ligand should also be provided. The approximate location of the binding site should also be specified, as RosettaLigand cannot perform binding site detection. Again, multiple parameters can be selected [33, 34]. Finally, two versions of AutoDock were also used, both tools that can be installed and run locally. AutoDock Vina was shown to be a strong competitor against six other programs when tested against a virtual screening benchmark [35]. The latest AutoDock software, AutoDockFR was also used.

AutoDockFR uses a genetic algorithm and scoring function based on the AutoDock4 scoring function. This program differs from the others as it takes into account receptor flexibility by allowing the user to specify flexible residues within the target structure, this allows the program to simulate induced fit caused by ligand binding where changes occur mainly in the residues side chains. This software was shown to outperform AutoDock Vina for tested datasets [36]. For both of the AutoDock versions, target and ligand structures are to be provided in PDBQT format. The search space, including the binding site, must also be specified with other optional parameters also available. AutoDock is commonly used for in-silico docking of associated drugs with HLA alleles. It is therefore important to assess the reliability of the different versions of this program when using the complex HLA structure examples.

Two drugs were investigated. Abacavir, an anti-retroviral drug used to supress HIV replication, is the most widely investigated drug associating ADRs with HLA. It has been shown that there is a genetic association between HLA-B*57:01 and Abacavir [4, 37, 38]. The ADR is thought to be driven by the activation of CD8+ T cells [39]. The mechanism of Abacavir binding has been experimentally validated by X-ray crystallography and shown to correspond with the altered peptide repertoire model. Abacavir binds directly and non-covalently with the HLA-B*57:01 binding cleft in the F-pocket (Fig 1) of the peptide-binding groove [40]. This binding results in an alteration of the physicochemical parameters and topography of the binding groove, altering the presented peptides and eliciting a polyclonal T-cell response leading to the Abacavir hypersensitivity reaction. Alterations at key residues within the binding cleft have been shown to prevent the Abacavir association, by testing closely related allotypes (e.g. HLA-B*57:03) and comparing the resulting hypersensitivity reactions seen in the risk B*57:01 allele [41].

The second drug investigated was Carbamazepine, an anti-epileptic drug, which has been strongly associated with Stevens-Johnson syndrome / toxic epidermal necrolysis (SJS/TEN), with patients having the B*15:02 allele showing hypersensitivity [5]. It is thought that the binding of Carbamazepine alters the self-presented peptides, through direct binding to the HLA molecule, similar to the Abacavir mechanism. These peptides are then recognised as foreign, leading to an immune response. Although the binding has not been experimentally validated, *in silico* modelling predicted the binding of Carbamazepine to HLA-B*15:02. It was predicted that the Carbamazepine binds in the D-pocket of B*15:02, adjacent to residue 156 of the HLA molecule [8]. This is one of the residues where the HLA-B*15:01 and HLA-B*15:02 alleles differ. As hypersensitivity is only seen in patients with the HLA-B*15:02 allele but not HLA-B*15:01, it is likely that this residue plays an important role in the ADR. A separate study has also predicted the binding site both through site-directed mutagenesis and *in silico* docking. The results of the site-directed mutagenesis implicated Asn63, Ile95 and Leu156, found in the D-pocket, in Carbamazepine presentation and T-cell activation as mutations at these positions (N63E, I95L or L156W) showed reduced binding affinity for Carbamazepine. *In silico* modelling conducted in the same study showed consistent binding near to the Arg62 residue located in the D-pocket of the peptide binding groove [14, 41]. These examples can therefore be used to test if the docking methods used predict the same binding position shown in these previous independent studies.

In this work, we used the Abacavir example for which a crystal structure of the complex exists, as a benchmark for the docking software. By using molecular docking, the binding position of the Abacavir within the B*57:01 risk allele HLA structure and, for comparison, with the non-risk control allele structures was predicted. Controls were chosen from B alleles shown to be non-risk (B*57:03) and common HLA-B and HLA-A alleles (B*07:02 and A*01:01). We work under the assumption that for (control) alleles that have not been associated with an ADR, that this is due to drug not binding sufficiently strongly to affect peptide presentation. Illing *et al*. showed that Abacavir interacts non-covalently with the B*57:01 risk allele but not with B*57:03 control [8]. The docking results were then compared to the known binding position to estimate the reliability of the docking protocol. In addition, we assessed to what extent the docking could distinguish between the HLA-associated and non-HLA-associated alleles. The same methods were used to test if the Carbamazepine binding position previously seen can be reliably replicated, using the programs showing the most accurate results for the Abacavir example. Due to there being more evidence available for the Abacavir example, including a crystal structure of the drug bound in complex, our investigations favour this example. For the Carbamazepine example, we are comparing our results against a previous prediction using similar methods.

This work sheds light on the utility and reliability of well-known docking programs used to address the challenging HLA-drug situation. These docking methods may help us to understand the mechanisms behind ADRs and identify genetic polymorphisms that may be influencing the binding seen in the risk but not control alleles.

## 2. Methods

### 2.1. Homology Modelling: Obtaining target structures

For the Abacavir example, B*57:01 has been shown to be a risk allele and B*57:03 not associated. For Carbamazepine B*15:02 was found to be the risk allele, with B*15:01 not being associated. These non-associated alleles were therefore used as controls along with a common HLA-B allele (B*07:02) and HLA-A allele (A*01:01) which could be assumed to not be associated as they are seen at a high frequency across European origin (Caucasian) populations (average frequencies obtained from AFND using gold standard populations [9]: B*57:01 = 0.03, B*07:02=0.10 and A*01:01=0.14).

The allele structures were obtained from the PDB database, where available. Models were made for those alleles where the structure is not publicly available (Table 1). Target and template sequences were aligned with ClustalX [42]. For each modelling exercise, ten models were generated using Modeller 9.9 automodel class and the model with the lowest objective function score was chosen as the model for docking. This objective function is a score generated from the spatial restraints and the CHARMM energy terms, reflecting stereochemistry within the structure [43].In these simple cases there was no need to explore alternative target-template alignments since sequences could be aligned unambiguously with no insertions or deletions.

**Table 1.** Structures of risk and control HLA alleles used in docking experiments. To distinguish between crystal structures and those created by homology models in the rest of the text, we add labels with suffix “sN” and “mN” where s means crystal structure and m means homology <u>m</u>odel, N is an integer where multiple models have been created.

| Drug | Risk Allele | Risk Allele Structure | Control Allele | Control Allele Structure |
| --- | --- | --- | --- | --- |
| Abacavir | B*57:01 | 3VRI [8] (*B5701_s*) | B*57:03 | 2BVP [44] (*B5703_s*) |
|  |  | Homology model using B*52:01 (3W39 [45]) and B*58:01 (5IM7 [46]) as templates (*B5701_m*) |  | Homology model using B*57:01 as a template (*B5703_m*) |
|  |  |  |  | Homology model using B*57:01 and B*07:02 (*B5703_m2*) |
|  |  | 2RFX [39] (*B5701_s2*) | B*07:02 | 4U1H [47] (*B0702_s*) |
|  |  |  | A*01:01 | 3BO8[48] (*A0101_s*) |
| Carbamazepine | B*15:02 | Homology model using B*15:01 | B*15:01 | 1XR9 [49] (*B1501_s*) |
|  |  | as template (*B1502_m*) | B*07:02 | 4U1H [47] (*B0702_s*) |
|  |  |  | A*01:01 | 3BO8 [48] (*A0101_s*) |

The Abacavir risk and control alleles were used to evaluate the homology modelling as the known structures are available for each allele investigated and so can be compared with the model structure and docking results. Two models were created for B*57:03, one with one template allele (B5703_m) and another with two template alleles (B5703_m2). These models could then be compared to the known structure of B*57:03 (B5703_s), as could the docking predictions, to understand the influence of these steps when employed in a typical docking protocol. The structure of B*57:01 was also modelled (B5701_m), from two similar sequences identified to make similar comparisons and evaluate the reliability of using homology modelling.

For the Carbamazepine risk associated allele B*15:02, there are four differing residues with control allele B*15:01. Three of these lie in the peptide binding groove, with only one of these being vital to the D-pocket architecture, where the Carbamazepine is predicted to bind (pos 156). Only a single template, the structure of B*15:01, was therefore used to model B*15:02 (B1502_m). The quality of each model was investigated using Ramachandran plots and QMEAN scores. For the B5701_m and B5703_ m and m2, RMSDs were used to give a measure of how well the models represent the known structures. The percentage sequence identity for each of the templates used for each model are shown in S1 Table.

#### 2.1.1. Conformational sampling of receptor protein structures

In order to identify the flexible side chains for the target structure, the relax function of Rosetta was used to explore the conformational properties of each residue. By looking at 10 different relaxed structures for each target, flexible residues can be identified. This allows us to consider the flexibility of the target structure when using AutoDockFR. When using the ROSIE server, a similar sampling is incorporated into the docking procedures [34].

### 2.2. Docking

#### 2.2.1. SwissDock

Using SwissDock [31], the default parameters search the whole target structure but by setting the search space parameters, it is possible to restrict binding to perform a local docking assay using the known binding pocket. Here, the area of interest is the peptide binding groove, including residues 1-180 of the alpha chain. The search space was therefore restricted to this area of interest. The file was processed to ensure it was in the correct format to be uploaded to SwissDock. This included removing the ligand and peptide from the structure. The PDB was then passed through the Prepdock server [50] to prepare the structure for docking. This prepared file was then submitted to SwissDock with the relevant known drug structures. SwissDock used the ZINC database to obtain the known structures of compounds (Abacavir ZINC ID: 2015928, Carbamazepine ZINC ID: 4785).

#### 2.2.2. >ROSIE

ROSIE [33, 34], was also used in a similar way. The PDB files were again prepared, removing the Abacavir and peptide from 3VRI and this time also removing the water molecules from the target structure, as Rosetta is unable to natively recognise these residues. The Abacavir drug structure was extracted from the relevant PDB (3VRI) and converted to SDF format. This was then submitted to the server to be docked, with the search space specified to centre on the peptide binding groove. This process was repeated for each of the risk and control alleles to be investigated.

#### 2.2.3. AutoDock

Two versions of AutoDock were also used, AutoDock Vina [35] and the later version AutoDockFR [36]. AutoDock Tools [51] was used to prepare the PDBQT files for both the target HLA alleles and the ligand structure. The drug structures for Abacavir was extracted from the 3VRI PDB file [8]. For Carbamazepine, there is no crystal structure available for the drug bound to a target and so the PDB file was obtained from the ZINC entry previously used for the SwissDock docking and from the Drugbank structure.

Using AutoDock Vina, a search space of 40×40×40Å was used. Initially, the default exhaustiveness was used but this was then increased gradually to identify the best parameters to find the closest docking poses of Abacavir to the native position seen in the crystal structure. Once this was identified, the process was then repeated for the other alleles, using the same parameters.

Using AutoDockFR, the docking was performed twice, either assuming the target as rigid or allowing for flexible residues. The aligned alleles and the ligand PDB files were converted to PDBQT files using AutoDock Tools [51]. The target structure and ligands were then loaded separately into AutoGridFR [52]. The pockets located within the target structure were identified and the search space selected to 1) surround the known binding position of the ligand, 2) surround the peptide binding groove and 3) surround the top three largest pockets identified. By selecting the top three pockets, this increases the search space and gives the opportunity for the ligand to bind in an alternative pocket and can tell us if the peptide binding groove is indeed the favoured binding region or not. The affinity maps are then generated by AutoGridFR and these are then inputted into AutoDockFR along with the ligand and the parameters for the docking. This process was repeated for each of the alleles, using the three different search spaces. As AutoDockFR only gives the binding pose for the top binding solution, AutoDockFR was ran in batches in order to obtain multiple binding poses. The top pose is therefore given for a selection of ten runs as opposed to the top ten poses for one run, as seen in the other examples.

Rosetta Relax was used to identify flexible side chains. These residues were then selected as flexible in AutoGridFR and the search space was set along the peptide binding groove, encompassing these residues. The affinity maps were generated and as before, inputted into AutoDockFR.

It was predicted that AutoDockFR may show more accurate docking predictions for unbound structures than those using the 3VRI crystal structure with both Abacavir and the peptide removed. As a result, the structure of HLA-B*57:01 without Abacavir bound (*B5701_s2*) was obtained from the PDB database (2RFX [39]). The peptide bound was removed and the B*57:01 structure was used in a similar way to give predicted binding when searching the peptide binding groove, assuming the receptor to be either rigid or flexible.

The results files from all the docking programs were then processed. The RMSDs (Root-Mean-Square Deviation) between the non-hydrogen atoms of the docked poses and the known binding position of Abacavir were calculated through PyMOL [30] using the “rms_cur” command. RMSDs were used to give a quantitative guide to how close the prediction poses lay to the known binding mode of Abacavir. The RMSDs along with visual inspection and docking scores were used to assess the reliability of each of the docking programs. The scores were also used to investigate the relationship between the binding scores and positions. The binding predictions for Carbamazepine were compared to predictions from previous studies in a similar way although RMSDs could not be measured for that case as no crystal structure was available.

## 3. Results

### 3.1. Homology Modelling

The first analysis was to explore the overall effect of homology modelling on the reliability of results from *in silico* docking. As expected, given the small number of amino-acid differences between models and templates, the QMean and Ramachandran plots for each of the alleles were shown to be within acceptable limits, showing the models to be of good quality. B5701_m was shown to be very similar to the known structure B5701_s with RMSD 0.81Å (266 to 266 atoms). The B*57:03 models also showed low RMSDs (B5703_m = 1.46Å (266 to 266 atoms), B5703_m2 = 1.24Å (266 to 266 atoms)) when compared to the known structure B5703_s. The two differing residues between the B*57:03 and B*57:01 alleles (position 114 and 116) both lie along the peptide binding groove and are vital for the architecture of the F-pocket shown to be the known binding position of Abacavir. It is therefore important that the model correctly represents the allele structure. When comparing the B*57:03 and B*07:02 control allele structures used for the Abacavir example with B5701_s, it could be seen that the tyrosine at position 116 for the B*57:03 and B*07:02 known structures overlaps with the known binding position of the Abacavir. It would be expected for the tyrosine in the B*57:03 model to show a similar conformation to that seen in these known structures. This was not seen in the B5703_m model but was seen in the B5703_m2 model in which two templates were used (S1 Fig). Using two templates for the model gave a more accurate representation of this element of the target structure in this case.

### 3.2. Abacavir

#### 3.2.1. All docking software assessed could dock Abacavir into the risk allele crystal structure but could not always predict the correct binding mode

Using SwissDock, it can be seen that the B5701_s example gives a docking solution close to the native position (Fig 2 a-c) with all poses showing binding in the F-pocket. When using AutoDockFR assuming the receptor to be rigid, similar results were seen to those using SwissDock (Fig 2 d-f). All poses for the B5701_s risk allele were shown bound in the same pocket as the known binding position. Little difference was seen between the docking shown for each search space, with only one pose from the B*57:01 run showing binding outside of the peptide binding groove when the search space was extended around the whole protein (data not shown). This indicates that the peptide binding groove is the most favourable region for binding. It can be concluded overall that the Abacavir docks in the expected binding pocket for these packages, but they cannot predict the exact native pose.

**Fig 2.**
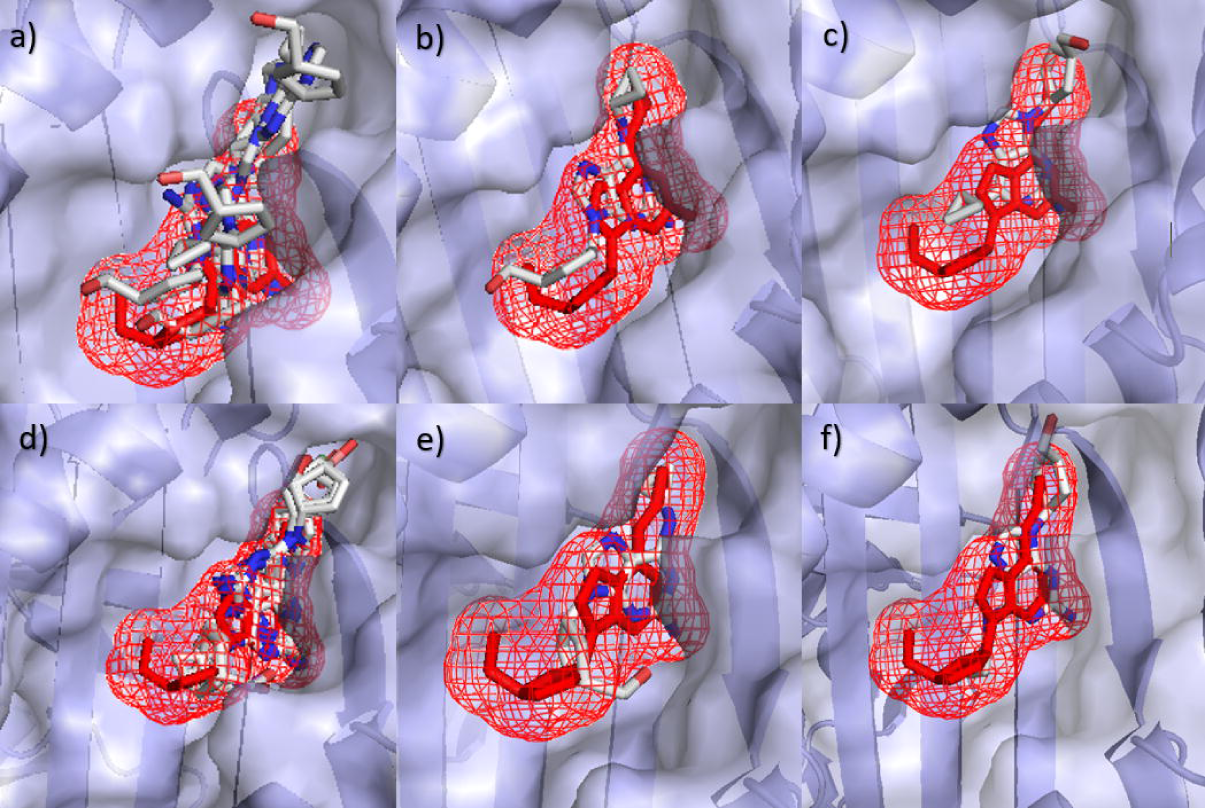
Binding positions for B*57:01 (B5701_s1) with Abacavir using SwissDock (a-c) and AutoDockFR (d-f). Predicted Abacavir binding positions using SwissDock (a-c) and AutoDockFR (d-f) are shown by white molecules with B5701_s1 shown as grey structure. The known binding position of Abacavir is shown in red. (a) All binding poses using SwissDock, (b) SwissDock pose 1 giving lowest RMSD of 2.02Å and (c) SwissDock pose 4 giving a higher RMSD of 6.92Å due to reversed orientation, (d) All binding poses using AutoDockFR, (e) AutoDockFR pose 3 giving lowest RMSD of 2.25Å and (f) AutoDockFR pose 5 giving a higher RMSD of 6.92Å due to reversed orientation. AutoDockFR docking ran using the search space centred on the known binding position of the ligand.

As the ROSIE server allows movement of side chains within the target, the modelled structure of the B*57:01 risk allele target was compared to the B5703_s structure submitted. This resulted in the RMSDs shown in Table 2 with B5701_s giving similar average RMSDs to the control alleles. Comparing the lowest RMSDs, the control structures give poses with lower RMSDs than B5701_s.

**Table 2.**
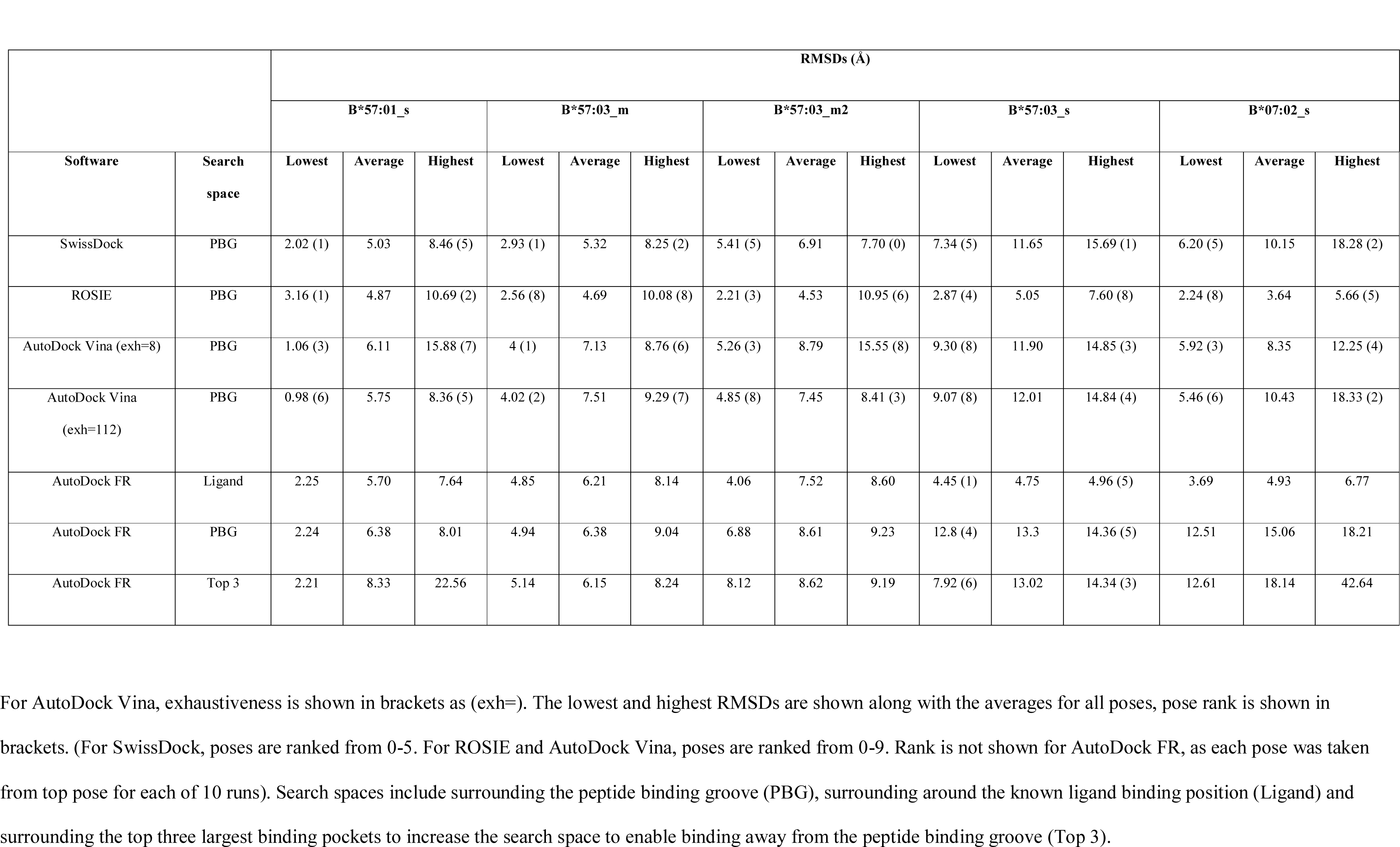
RMSD measurements for Abacavir docked with B*57:01, B*07:02 and B*57:03 models, using the different software.

When docking the Abacavir structure using AutoDock Vina, the exhaustiveness was investigated using the B*57:01 example (B5701_s) to optimise the protocol before docking the other non-risk alleles. Using the default exhaustiveness of 8, the RMSD values were shown to be quite variable (Table 2). The exhaustiveness was then increased starting at exh=18 and doubling the exhaustiveness to see the effect. It was found that the exhaustiveness of 112 gave the lowest RMSDs overall and these did not improve with further increasing of exhaustiveness

#### 3.2.2. Most docking software assessed can distinguish between risk and control alleles

Two methods were used to investigate if the docking software can distinguish between the risk alleles. RMSD values, alongside some visual inspection, were used to give a measure of how similar the docking prediction results are to the known binding position obtained from the crystal structure. Docking scores were also investigated to see if there is more favourable binding seen in the risk alleles compared to the controls and also how the scores differ between the predicted poses for the risk allele itself.

Using RMSDs, it can be seen from Figure 3 and Table 2 that most of the software, excluding “AutoDockFR (Ligand)” and ROSIE, showed lower RMSD’s for B*57:01 than the other control alleles investigated and were able to distinguish between the known risk allele structure (B5701_s) and the known control allele structures (B0702_s and B5703_s). AutoDockFR using a search space surrounding the known binding position of the Abacavir ligand, “AutoDockFR (Ligand)”, shows lowest median RMSD’s for B*07:02. ROSIE shows lowest median RMSD’s for all the alleles investigated. AutoDock Vina (with exhaustiveness 112) was able to achieve the lowest RMSD overall for B*57:01 (0.98 Å) but showed more variability between poses, giving a higher average RMSD, although this was still lower than the average RMSDs for the control alleles. Using SwissDock and AutoDockFR, the control alleles showed higher RMSDs than the risk B*57:01 allele although it can be seen that some poses gave higher RMSDs, similar to those seen for the control alleles. Some of the predicted poses for the B*57:01 allele were shown to be binding in the correct pocket but gave a higher RMSD due to the reversed orientation (Fig 2c & 2f). By examining these poses it can be seen that the ligand makes similar interactions to the correct binding pose (S2 Fig) but the reversed orientation results in the higher RMSD. It is therefore important to consider both the poses and the RMSD scores when comparing binding results.

**Fig 3.**
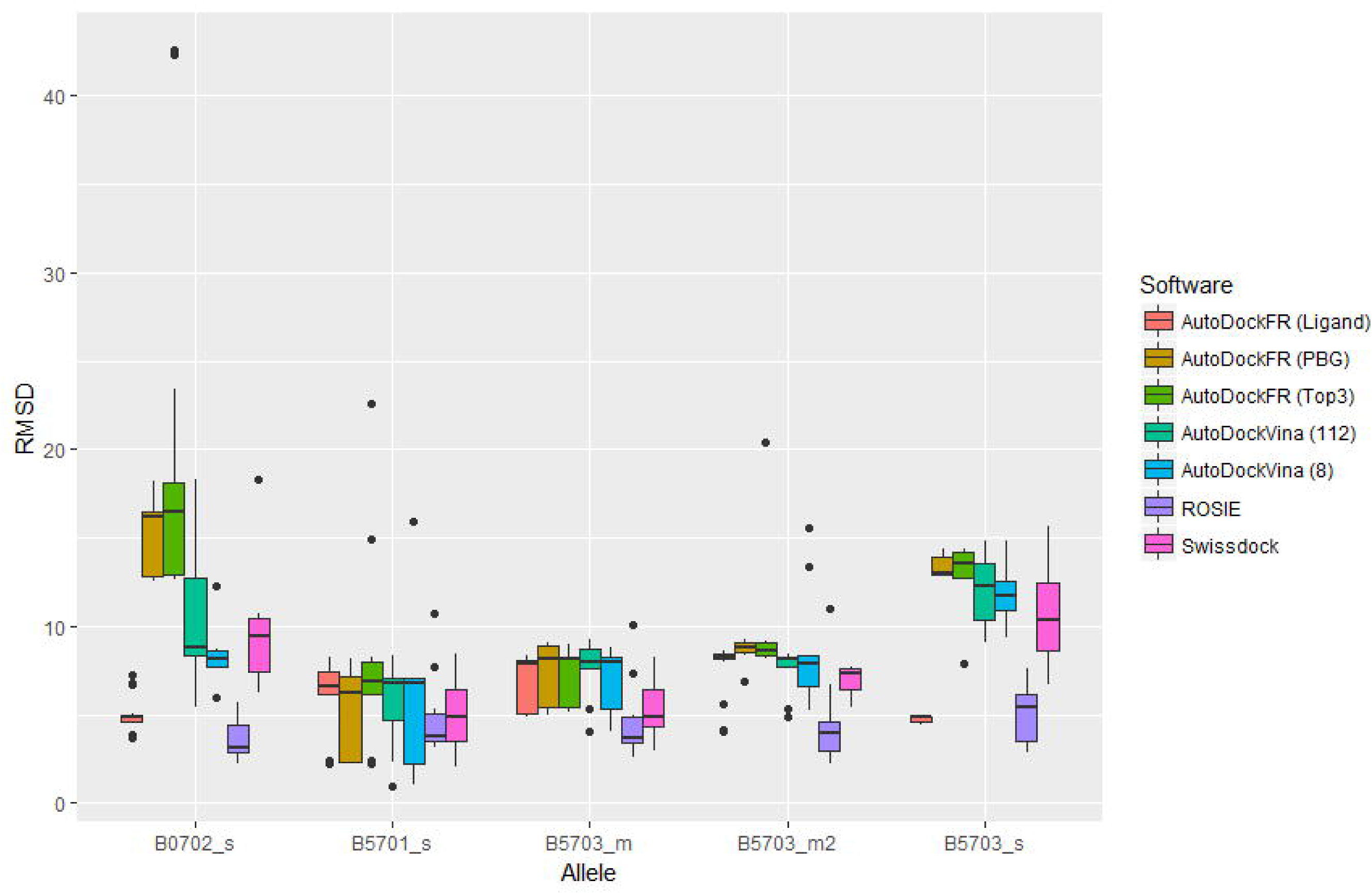
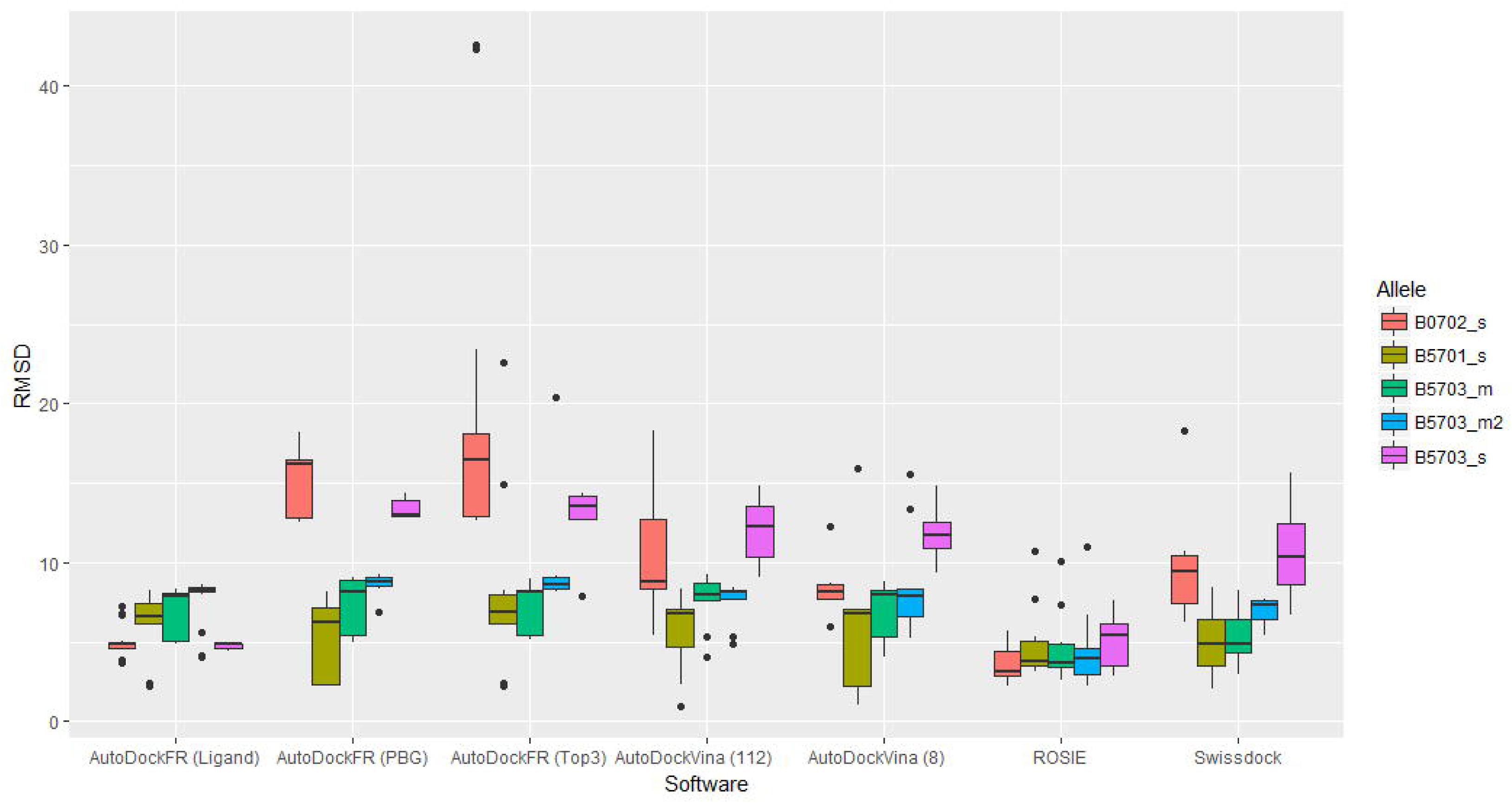
Boxplots to show RMSD values of Abacavir poses with respect to the known structure. Plots grouped by (a) Allele and (b) software. Search spaces for AutoDockFR are shown in brackets and include surrounding the peptide binding groove (PBG), surrounding around the known ligand binding position (Ligand) and surrounding the top three largest binding pockets to increase the search space to enable binding away from the peptide binding groove (Top 3). For AutoDock Vina, exhaustiveness is shown in brackets.

Figure 4 shows the docking poses for the control alleles using SwissDock. It can be seen that the Abacavir binds further from the known binding position from 3VRI, with A*01:01 showing the largest difference from the B*57:01 allele, predicting binding away from the peptide binding groove (RMSDs: lowest = 21.44 Å, average = 23.23 Å, highest = 25.47 Å).

**Fig 4.**
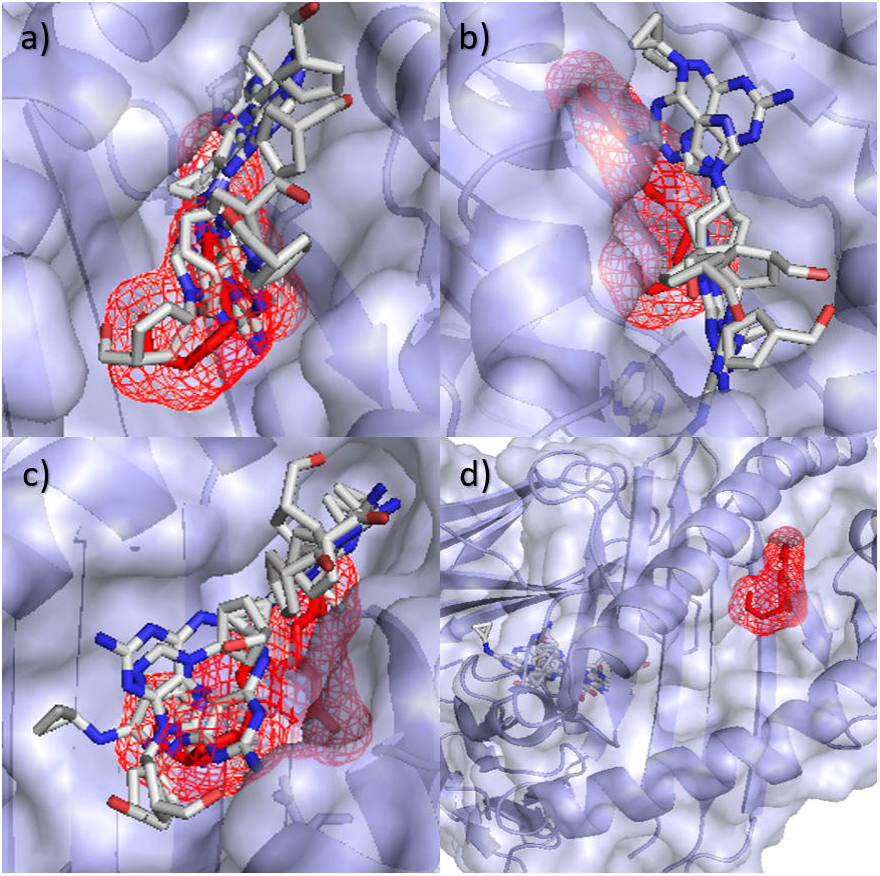
All docking poses of Abacavir for control alleles using SwissDock. (a) B*57:03 using one template (*B5703_m*), (b) B*57:03 using two templates (*B5703_m2*), (c) B*07:02 (*B0702_s*) and (d) A*01:01 (*A0101_s*). Known binding position of Abacavir shown in red. Poses can be seen further from the native pose than found in the risk allele docking.

It can be seen that B*57:01 generally had a lower average RMSD than the control alleles for all software, excluding ROSIE, even with these reversed orientations discussed, with the lowest RMSD being constantly lower than those seen for the controls. Both control alleles B*57:03 and B*07:02 contain a tyrosine at position 116, rather than the serine seen in the risk allele, with A*01:01 having an aspartic acid at position 116. This residue is sensitive for the F-pocket architecture as it lies along the base of the pocket [53] and so this mutation prevents binding in this native position.

The full fitness scores for SwissDock poses were investigated, with lower scoring poses being more favourable than higher scoring poses. Scores were compared between B*57:01, B*57:03 and B*07:02, it was found that poses for the B*57:01 risk allele scored more poorly than those for the non-risk alleles (Fig 5a), with the control alleles showing lower scores than the risk allele. This implies that comparison of scores between alleles is not valid since better docking results were seen for the risk allele. Put another way, the docking scores were not able to distinguish between the risk and control alleles. Nevertheless, the scores for the B*57:01 risk allele are a good guide to pose accuracy, as there is a modest positive correlation between RMSD and score (Fig 5b), with an R^2^ value of 0.65.

**Fig 5.**
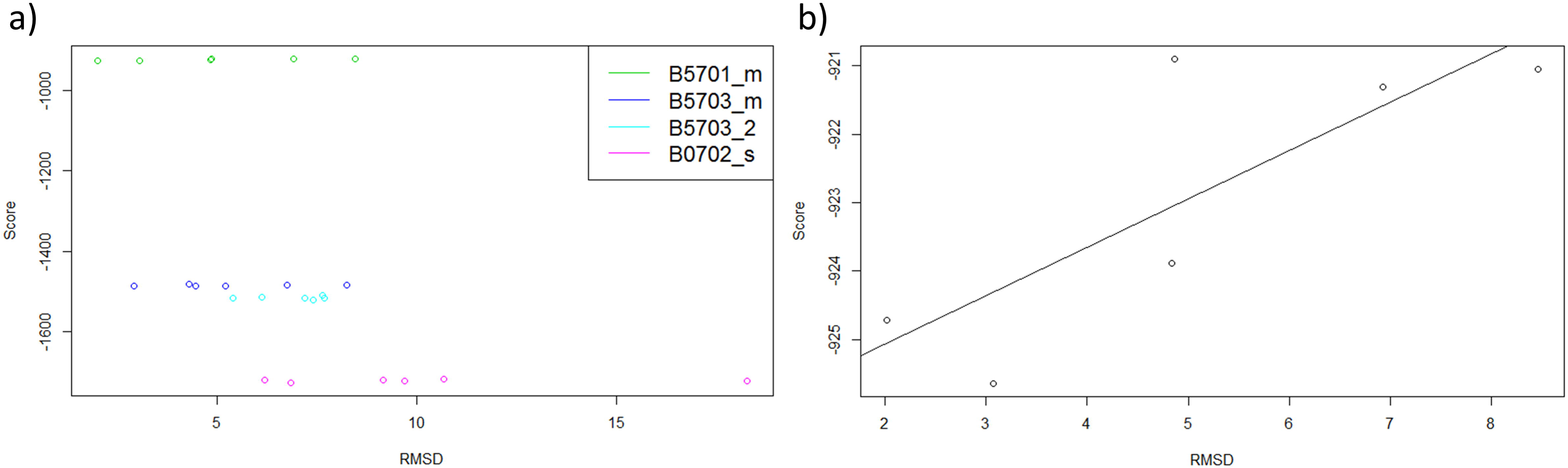
Full fitness scores versus RMSD. (a) Scatterplot to show the full fitness scores vs RMSDs for each pose for each of the different alleles. The non-risk allele poses have lower scores than poses for the risk allele. (b) Scatterplot to show full fitness score vs RMSD for B*57:01 poses.

#### 3.2.3. Docking performance can be degraded by using a homology model

The docking of Abacavir with the known and modelled structures of B*57:01 and B*57:03 were compared (S3 Fig). This allowed us to compare the docking positions between known and modelled structures. B5701_m showed an unexpected overlap between the Ser116 residue and the known binding position on Abacavir from 3VRI. This prevented the docking from predicting the exact native pose. However, poses were still predicted within the F-pocket and had lower RMSDs seen than those predicted for the non-risk alleles. This slight difference between the modelled and known structures may be due to the peptide bound in the peptide binding groove of the 3VRI structure.

The docking of Abacavir to the known structure of the control allele B5703_s showed similar results to the modelled structures, with poses seen away from the known Abacavir binding position and higher RMSDs (S4 Fig). The average RMSDs seen with the B*57:03 known structure were higher than those seen with the homology models, showing the Abacavir docks further from the known binding position with the known structure.

#### 3.2.4. Receptor flexibility negatively affects the docking performance

When the flexible residues were incorporated for the 3VRI docking with Abacavir example, poses occupied the whole peptide binding groove and did not favour the F-pocket as expected (Fig 6). When the scores for the poses found inside the F-pocket were compared to those found outside the F-pocket, it was also seen that these scores did not favour the F-pocket (data not shown). Thus, although the complex algorithms developed for AutoDockFR have been shown to improve the success of docking [36] in general, they degraded performance in our example, suggesting that they should only be used with caution for HLA docking.

**Fig 6.**
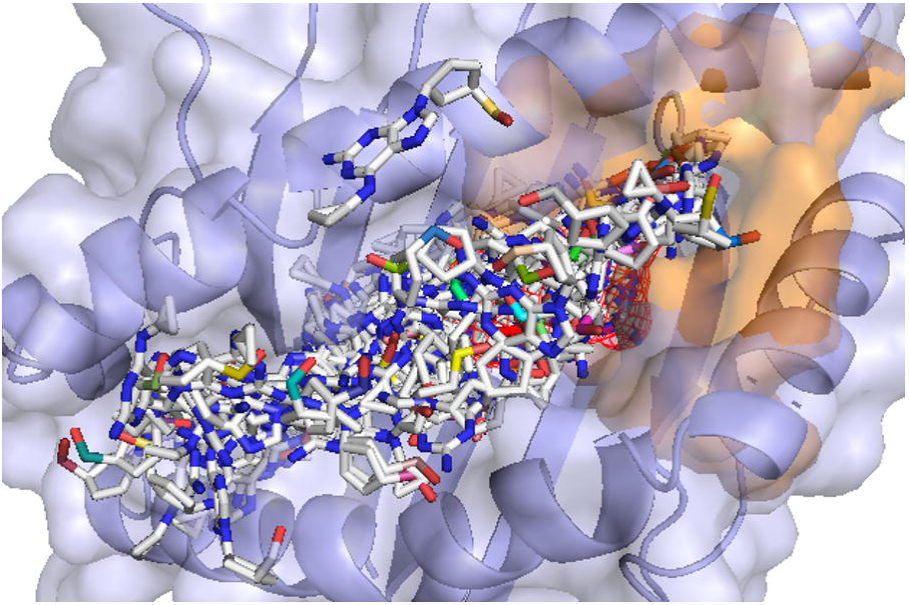
Abacavir docked with B*57:01 structure (3VRI), using ADFR assuming the receptor to be flexible. Poses are seen along the whole groove and not just the F-pocket (shown in orange). The docking is unable to predict the native-like poses (known binding of Abacavir shown in red).

#### 3.2.5. Using AutoDockFR cannot compensate for the added difficulty of docking to the unbound target

AutoDockFR was also used to dock Abacavir with the unbound structure of HLA-B*57:01 (2RFX [39]), crystallised in the absence of drug, in order to test the possibility that the structure without Abacavir bound would yield better docking results when flexibility of residues was considered. The peptide was removed from 2RFX and the B*57:01 structure was used as the target. Again, two runs were completed, assuming the receptor to be either rigid or flexible.

Docking to the structure assuming rigid side chains produced poses in both the B and F pockets (Fig 7a). However, the lowest RMSD is seen as 8Å and is therefore not very accurate when comparing to the known binding position. When looking at the poses, it can be seen that the native pose could not be predicted. When flexible residues were incorporated with AutoDockFR (Fig 7b), the entire peptide binding groove was again occupied.

**Fig 7.**
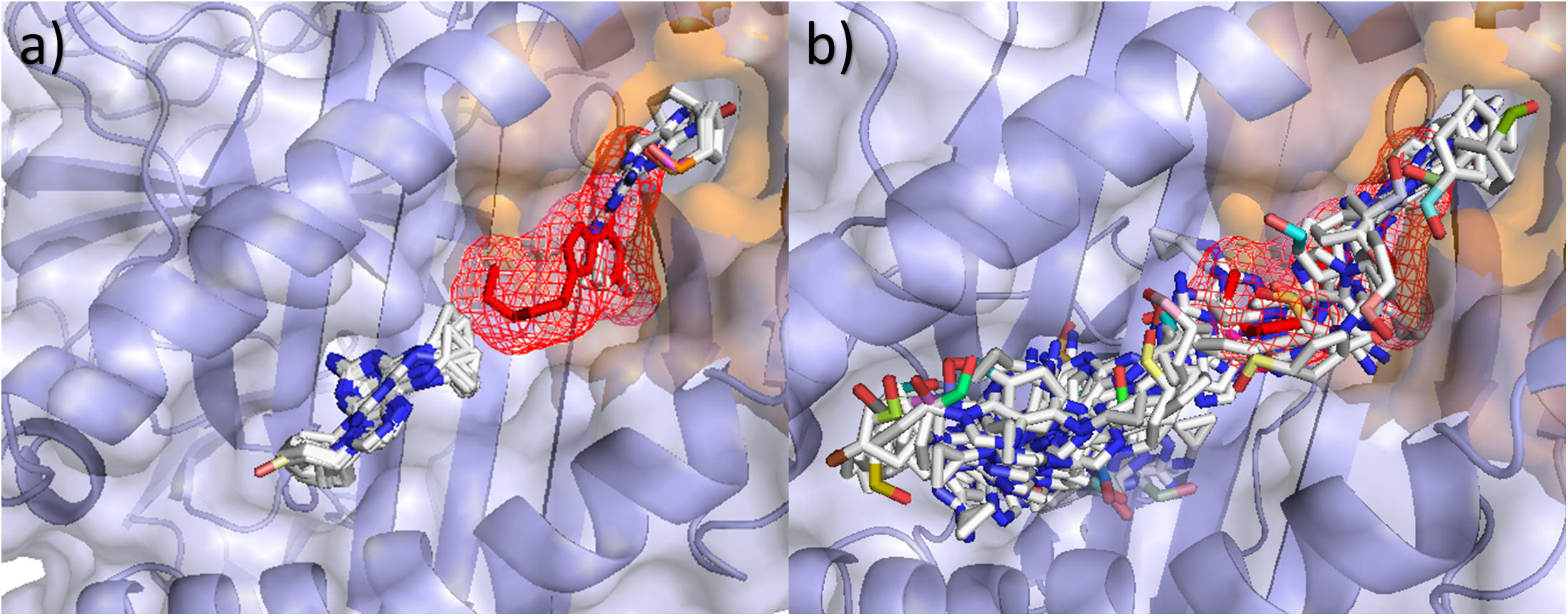
Abacavir docked with unbound B*57:01 structure (2RFX), using ADFR. (a) Assuming the receptor to be rigid. Only a few poses can be seen bound within the F-pocket (shown in orange), with these poses showing incorrect conformation. The docking was unable to predict native-like poses (known binding of Abacavir shown in red). (b) Assuming the receptor to be flexible. Poses are found along the whole groove and not just the F-pocket (shown in orange). The docking is unable to predict the native-like poses (known binding of Abacavir shown in red).

Although AutoDockFR gives good results for the ideal case when docking the Abacavir structure back into the B*57:01 structure obtained from 3VRI (by removing both the ligand and the peptide), it is shown here that docking using the unbound structure, crystallised in the absence of the drug, showed less accurate results and the correct binding position could not be identified. This highlights the difficulties of using docking to investigate these challenging HLA-ADR cases as in general docking for new ADRs will be performed, for example, on structures that have a peptide already bound but not the drug that is to be docked.

### 3.3. Carbamazepine

SwissDock and AutoDockFR were best able to predict the binding positions of Abacavir and so these programs were both used to predict the binding position of Carbamazepine with both the risk allele (B1502_m) and the control alleles (B1501_s, A0101_s and B070_s2). The predicted poses were compared to those shown in previous studies in which B*15:02 showed binding in the D-pocket, close to residues 62, 63, 95 and 156 [8, 14].

Docking Carbamazepine with B1502_m using SwissDock, the poses were predicted to sit in the D-pocket previously identified as of interest. Looking at the docking results (Fig 8a), it can be seen that Carbamazepine is predicted to bind in the D-pocket of B*15:02, close to Leu156, identified by the study as important, with only one pose predicted out of this pocket. Using AutoDockFR, the same general pattern was seen with the D-pocket generally being favoured for B*15:02 (Fig 8b). Using SwissDock, the B*15:01 docking showed poses predicted to bind elsewhere, away from this pocket, as predicted. For the B*15:01, A*01:01 and B*07:02 alleles, with a mutation at this 156 position (Leu→Trp, Leu→Arg and Leu→Arg respectively), this D-pocket is closed off and produces poses elsewhere (S5 Fig). Using AutoDockFR, the same general pattern was seen with no poses being found in or around the D-pocket for B*15:01. These predictions cannot be validated since there is, as of yet, no crystal structure available for Carbamazepine bound to HLA. However, when compared to previous studies, our results showed a similar binding position using different software.

**Fig 8.**
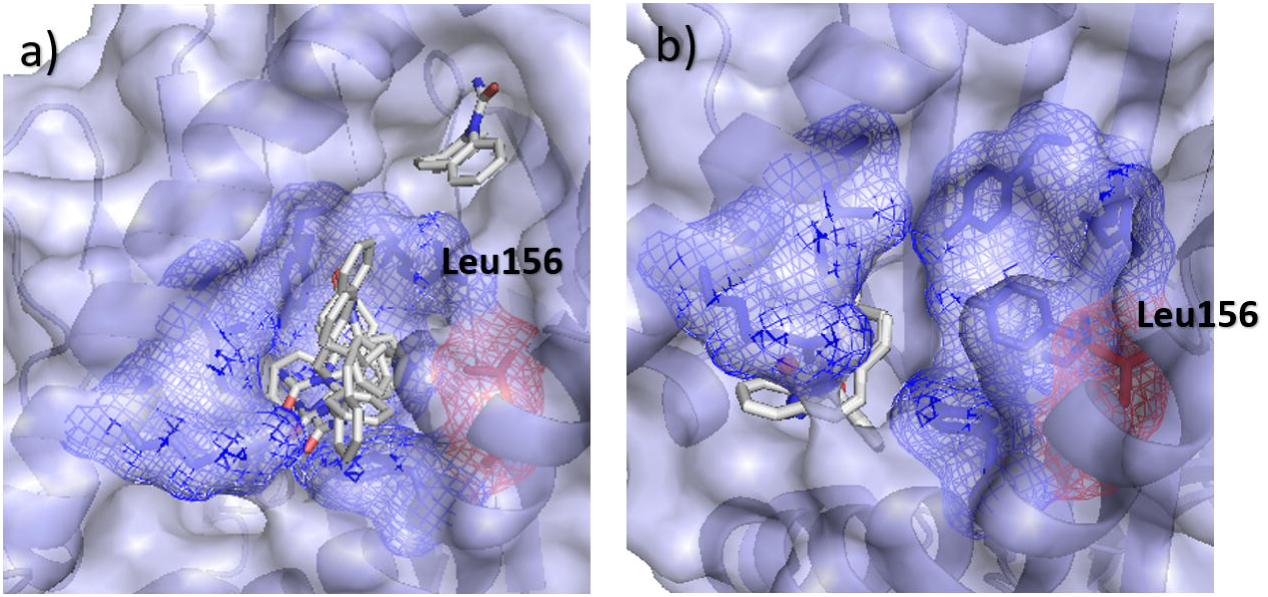
Docking Carbamazepine with B*15:02 risk allele using SwissDock and AutoDockFR. (a) SwissDock and (b) AutoDockFR poses for Carbamazepine docked with B*15:02 risk allele. Residue at 116 shown in red, with other D-pocket residues shown in blue.

## 4. Discussion

The purpose of this exercise was to compare multiple docking programs to assess their performance with these challenging HLA-ADR cases. We used the Abacavir example as the benchmark for the docking software, as the crystal structure is available for Abacavir bound in complex with its associated risk allele. We used the different docking programs to re-dock the Abacavir into the risk allele and measure how accurately each program could predict the known binding position. It was found that the binding modes can sometimes be predicted but not always. Each docking program used was able to predict the Abacavir binding within the F-pocket and, with the exception of the ROSIE server, was also able to distinguish between the risk and control alleles with the best scoring poses for the control alleles being seen further from the known binding position and in some cases, away from the F-pocket. This was also generally reflected in the RMSDs, although it is important to consider these alongside the poses themselves as reversed orientations can give higher RMSDs for poses found close to the known binding position.

Using the same docking methods to investigate the Carbamazepine example, the docking programs used were able to recreate previously published predictions. The docking software was also able to distinguish between the risk and control alleles with the risk allele showing binding in the D-pocket and the other control alleles showing poses away from the D-pocket.

Here we also investigated the impact of homology modelling on the docking performance. Homology models are commonly used for docking, especially with *in silico* database screening and have been shown to give accurate docking predictions [54–57]. However, localised errors can still have a big impact and so special caution should be used when there is no crystal structure available for the docking. It is important to ensure the models are as accurate as possible in order to give accurate predictions. Especially when mutations may impact the architecture of the predicted binding pocket, such as in the Abacavir example.

Predictions of docking poses have subsequently been experimentally validated by a crystal structure. Yang et al. [58] predicted binding of Abacavir to B*57:01 in the F-pocket and predicted positions 114 and 116 as important for binding. This was then confirmed once the crystal structure of the HLA-drug complex was determined [8]. Other predictions have also been made which fit well with experimental data, about binding of Carbamazepine, for example, and have been in accord with the experimental data [5].

Caution is needed to overcome the challenges produced from docking with the HLA structures. The long hydrophobic peptide-binding cleft is separated into subsites and small molecules can bind in any of these subsites along the cleft. The binding of drugs to HLA is probably weaker than many natural and drug-target interactions as the HLA binding site has not naturally evolved to recognise the drug, nor has the drug been designed or discovered in a structure-based fashion. This presents challenges for docking as there may be fewer interactions formed and less steric complementarity between the drug and its HLA recognition site. A further complication is the possibility that bound peptides may stabilise drug poses that would not otherwise be energetically favourable. Addressing this issue computationally is beyond the capability of current tools but the possibility should be borne in mind.

Furthering our understanding of the potentials and limitations of docking small molecules to HLA is important to aid our understanding of the underlying mechanisms involved with these ADRs.

Understanding these mechanisms and how the binding of small molecules varies between risk and control alleles may enable us to make predictions of potential ADRs by identifying polymorphisms which may contribute to direct binding. This may also lead to improved understanding and predictions of ADRs, ultimately leading to reduced risk due to screening procedures.

## Acknowledgements

The research was part-funded by the Medical Research Council grant for the Centre for Drug Safety Science, University of Liverpool (Grant Number: MR/L006758/1). The funders had no role in study design, data collection and analysis, decision to publish, or preparation of the manuscript.

## Supporting information

**S1 Table. Percentage sequence identity between models and templates.** Table to show the percentage sequence identity calculated using BLAST-P [59] for each of the template sequences compared to the model sequences.

**S1 Fig. Crystal structure of B*57:03 aligned to modelled structure of B*57:03.** Crystal structure of B*57:03 (green) shown aligned with models created using one template (blue) and two templates (pink). Also shown with the known binding position of Abacavir (red). Looking at Tyr116 shown highlighted as sticks, it can be seen that the B*57:03 model created using one template shows a different conformation than expected by the known structure.

**S2 Fig. Ligplot plots show the interactions between Abacavir and B*57:01.** (a) Known binding position of Abacavir in complex with B*57:01 (3VRI). (b) B5701_s using AutoDockFR, pose 3 showing lowest RMSD (2.254 Å). (c) B5701_s using AutoDockFR, pose 4 showing highest RMSD (7.639 Å). Similar interactions with key residues can be seen between all poses (circled). Dashed lines show Hydrogen bonds (with length), spoked arcs show hydrophobic bonds. Created using Ligplot [56].

**S3 Fig. Comparison of docking poses using crystal and modelled structures of B*57:01 and B*57:03.** (a) Known structure of B*57:01 (B5701_s) showing all docking poses for Abacavir using SwissDock; (b) Modelled structure of B*57:01 risk allele (B5701_m) showing all docking poses for Abacavir using SwissDock; (c); B*57:03 known structure (B5703_s) showing all docking poses for Abacavir using SwissDock; (d) modelled structure of B*57:03 (B5703_m2) showing all docking poses for Abacavir using SwissDock. Known binding position of Abacavir from 3VRI shown as red mesh.

**S4 Fig. Comparison of RMSDs for docking poses using crystal and modelled structures of B*57:01 and B*57:03.** Boxplot to compare the RMSDs for poses compared to the known binding position of Abacavir, for both the crystal structures and models of B*57:01 and B*57:03 using SwissDock.

**S5 Fig. Docking Carbamazepine with control alleles using SwissDock.** SwissDock poses for Carbamazepine docked with (a) B*15:01 control allele (B1501_s), (b) B*07:02 control allele (B0702_s) and (c) A*01:01 control allele (A0101_s). Residue at 116 shown in red, with other D-pocket residues shown in blue.

